# Impact of the MX segment on the biogenesis of *α*7 nACh receptors

**DOI:** 10.64898/2026.04.02.715926

**Authors:** Quynh Hoa Do, Irina Kim Cavdar, Petar N. Grozdanov, Joshua J. Theriot, Rhea Ramani, Michaela Jansen

**Affiliations:** Cell Physiology and Molecular Biophysics, School of Medicine, Texas Tech University Health Sciences Center, 3601 4^th^ Street, Lubbock, TX 79430, USA; Pharmacology and Neuroscience, School of Medicine, Texas Tech University Health Sciences Center, 3601 4^th^ Street, Lubbock, TX 79430, USA

## Abstract

Nicotinic acetylcholine receptors (nAChRs) belong to the pentameric ligand-gated ion channel superfamily (pLGICs). Among them, the neuronal homomeric α7 nAChR is highly permeable to calcium and plays critical roles in synaptic transmission, cell signaling, and inflammation modulation. The biogenesis of α7 nAChRs is enhanced by the chaperone proteins RIC-3 and NACHO. Previously, we reported a motif in the 5-HT_3A_ receptor, another pLGIC, involved in RIC-3 modulation. Residues in this motif are conserved and also found within the L1-MX segment of the α7 nACh subunit. We therefore explored the regulatory roles of these conserved residues in the biogenesis of α7 nAChRs using multiple approaches, including heterologous expression in *Xenopus laevis* oocytes, mutagenesis, pull-down assays, cell-surface labeling, and two-electrode voltage-clamp (TEVC) recordings. We find that synthetic α7 L1-MX peptide interacts with both RIC-3 and NACHO. In particular, conserved residues W330, R332, and L336 in the L1-MX positively regulates the assembly of α7 oligomers and the biogenesis of α7nAChR. In presence of residues W330, R332, and L336, NACHO promotes an assembly of an α7 pentamer which is resistant to strong denaturing conditions. NACHO-promoted α7 pentamer is also resistant to Endo H enzyme. Sensitivity of the pentamer to moderate temperatures (37 °C, 45 °C, and 50 °C) suggests that NACHO stabilizes the pentamer via non-covalent interactions. In contrast, Ala replacements at these residues disrupt the biogenesis and abolish α7 current. NACHO and RIC-3 co-expression yields partial rescue of functional expression for some Ala replacement constructs.

**SUMMARY:** This work identifies regulatory roles of conserved residues W330, R332, and L336 in the biogenesis of α7 nAChR. This discovery positions MX subdomain as a promising target for future drug development that can minimize adverse effects.

## INTRODUCTION

Neuronal nicotinic acetyl choline receptors (nAChRs) are assembled from various alpha (α2-α10) and beta (β2-β4) subunits to form diverse receptor subtypes (Paterson and Nordberg, 2000; Millar, 2003; Zoli et al., 2015). Among them, α7 nAChRs are highly permeable to calcium and plays critical roles in synaptic transmission, synaptic plasticity, and neuroprotection (Bertrand et al., 1993; Broide and Leslie, 1999; Jones et al., 1999; Khiroug et al., 2003; Wang et al., 2003; Fucile, 2004; Shen and Yakel, 2009; Wu et al., 2021). Consequently, dysregulations in the biogenesis of α7 nAChR are linked to multiple disorders, including Alzheimer’s disease, schizophrenia, autism disorder spectrum, and cancers (Whitehouse et al., 1988; Freedman et al., 2000; Guan et al., 2000; Lee et al., 2002; Davis et al., 2009; Hegde and Keenan, 2022; Terry et al., 2023).

α7 nAChR belongs to the superfamily of pentameric ligand-gated ion channels (pLGICs), and these receptors share a similar overall molecular structure, as shown in Fig. 1 A and B (Karlin, 2002; Lester et al., 2004; Dani and Bertrand, 2007; Albuquerque et al., 2009; Noviello et al., 2021). Each channel is formed by five individual subunits arranged around a central pore. Each subunit consists of an extracellular domain, a transmembrane domain (TMD) with four α-helical segments (M1 through M4), and an intracellular domain (ICD). The ICD, which connects the M3 and M4 segments of the TMD, or the M3-M4 loop, comprises an L1 loop, an MX helix, another large loop (not shown here), and the MA segment (Fig. 1 B).

**Figure 1.**
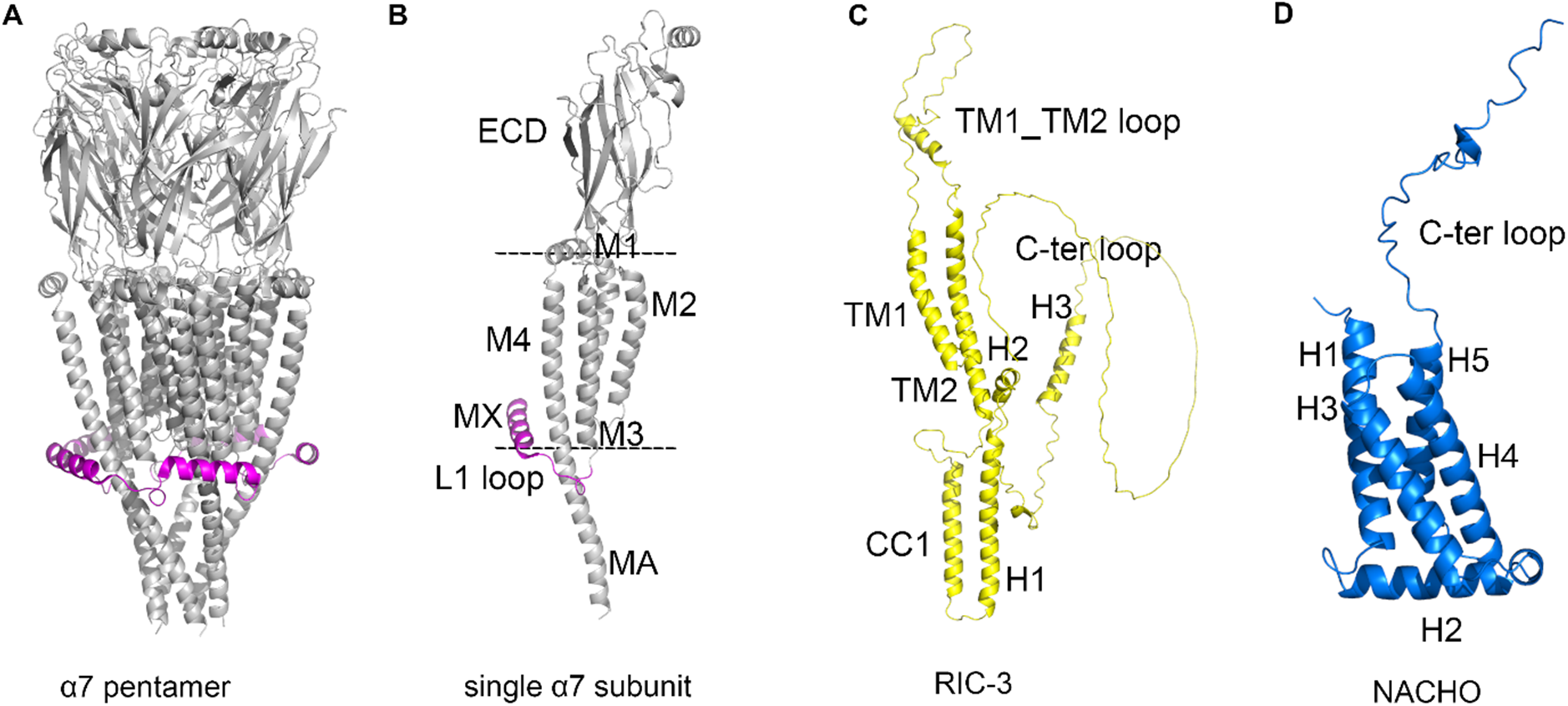
Structures of α7 nAChR, RIC-3, and NACHO. Ribbon representations viewed from the plane of the membrane. **A,** Pentameric channel assembly of α7 nACh subunits (PDB ID 7KOO), and **B**, a single α7 nACh subunit. The dashed lines illustrate the membrane limits. The extracellular domain (ECD) and four α-helical segments (M1 through M4) of the transmembrane domain are presented. For the intracellular domain connecting M3-M4, we show the L1-MX fragment (magenta) and the MA helix. The large loop connecting L1-MX and MA is not shown. **C,** The RIC-3 structure was predicted using Alpha Fold 3 and the amino acid sequence in UniProt Q7Z5B4. Consistent with previous studies, RIC-3 is predicted to contain two alpha-helical segments that have been described as transmembrane segments (TM1 and TM2) and an alpha-helical segment that represents the canonical coiled-coil domains (CC1) (Halevi et al., 2003; Treinin, 2008). H1, H2, and H3 are distinct helical segments following the CC1 domain. **D**, The NACHO structure was predicted using Alpha Fold 3 and the amino acid sequence in UniProt Q53FP2. Both N- and C-termini are located on the same side of the membrane plane. The N-terminal is followed by an α-helical segment H1 spanning the membrane. Following H1 is an α-helical segment (H2) parallel to the membrane plane. After H2 are three α-helical segments spanning the membrane (H3, H4, H5). And after the H5 helix is the C-terminal loop.

Previous studies have revealed that various proteins, including scaffold, regulatory, anti-apoptotic, and chaperone proteins, regulate the surface expression of α7 nAChR. Among them, PICK1 and Ly6h downregulate the functional expression of α7 nAChR’s, while anti-apoptotic Bcl-2 proteins, RIC-3 and NACHO upregulate this expression (Lansdell et al., 2005; Williams et al., 2005; Baer et al., 2007; Puddifoot et al., 2015; Gu et al., 2016; Dawe et al., 2019).

RIC-3, encoded by the *ric-3* gene (<u>r</u>esistance to inhibitors of <u>c</u>holinesterase), regulates the functional surface expression of pLGICs, including nACh and 5-HT_3A_ receptors (Halevi et al., 2002; Castillo et al., 2005). RIC-3 is expressed in various species, and in humans, is found in multiple tissues, including the immune and central nervous systems (Wang et al., 2009; Ben-David et al., 2020). The three-dimensional atomic structure of RIC-3 has not yet been experimentally determined. Using AlphaFold 3, we generated a predicted structure of human RIC-3 as shown in Fig. 1 C (Halevi et al., 2003; Abramson et al., 2024).

NACHO, encoded by the *tmem35a* gene, is a membrane protein and an endoplasmic reticulum resident, whose three-dimensional atomic structure has also not been experimentally determined (Tran et al., 2010; Kennedy et al., 2016). Using AlphaFold 3, we generated a predicted structure of NACHO, shown in Fig. 1 D. Previous studies have demonstrated that the M2 segment of the α7 subunit is critical for NACHO-mediated assembly of the α7 nAChRs (Kweon et al., 2020).

Our previous work demonstrates that RIC-3 regulates the surface expression of 5-HT_3A_ through direct interaction with a binding motif, D**W**L**R**…V**L**DR (Jansen et al., 2008; Nishtala et al., 2016; Pirayesh et al., 2020; Do and Jansen, 2023). The conserved residues in this motif (Trp, Arg, and Leu) are also present in the L1-MX segment of the ICD of the nACh α7 subunit (Fig. 2 A). RIC-3 can regulate the surface expression of both α7 nACh and 5-HT_3A_ receptors (Castillo et al., 2005). We, therefore, characterized roles of conserved residues, W330, R332, and L336 of the α7 L1-MX segment in regulating the biogenesis of α7nAChRs using multiple approaches, including pull-down assay, mutagenesis, expression of full-length α7 subunits in absence and presence of RIC-3 and / or NACHO in *Xenopus laevis* oocytes, cell surface labelling, and two-electrode voltage-clamp (TEVC) recordings.

**Figure 2.**
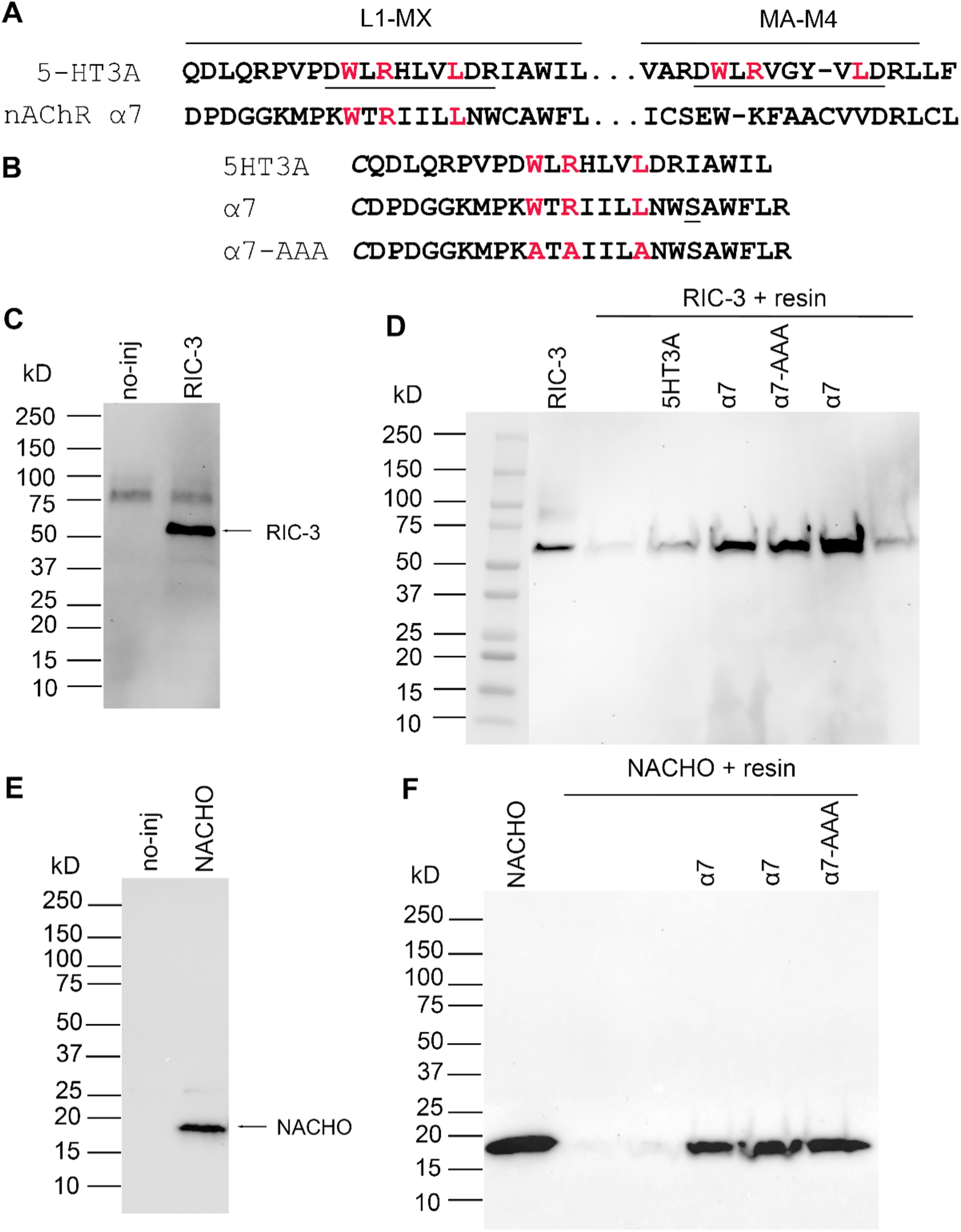
L1-MX segment derived from *α*7 nACh subunit interacts with RIC-3 and NACHO. Ala substitutions at conserved sites in L1-MX peptide retain the interactions with RIC-3 and NACHO. **A,** Sequence alignment of L1-MX segment and MA-M4 boundary from 5-HT_3A_ (UniProt Q8K1F4) and α7 nACh (UniProt P49582) subunits. Conserved residues Trp, Arg, and Leu in the RIC-3 binding motif (underlined residues) in L1-MX and MA-M4 in the 5-HT_3A_ subunit are highlighted in red. **B,** L1-MX peptides designed based on L1-MX segments of 5-HT_3A_ (UniProt Q8K1F4) and α7 nACh (UniProt P49582, C339S) subunits. Added N-terminal Cys (*C*) is italicized, and α7 Cys339Ser substitution (<u>S</u>) is underlined. **C,** Western blot image of RIC-3 protein extracted from *Xenopus laevis* oocytes not injected (no-inj) and injected (RIC-3) with RIC-3 IVT mRNA. In RIC-3 injected oocyte sample, a prominent band just above 50 kDa is detected. **D,** Western blot image of RIC-3 extracted from *Xenopus laevis* oocytes and RIC-3 in eluates from pull-down assays using 50 mM L-cysteine-capped resin and 3A, α7, or α7-A peptide covalently-linked resin. The eluates from peptide-linked resins give rise to stronger RIC-3 signals as compared to the eluate from the capped resin. Peptide/resin ratios in pull-down assays are 7.0 mg/ml for 3A, 8.8 mg/ml for or 6.7 mg/ml α7, and 5.9 mg/ml for α7-A. Data shown here are representatives of at least three biological replicates. Biological replicates are pull-down assays conducted using different RIC-3 extracts from various batches, each containing 70-110 *Xenopus laevis* oocytes. Each biological replicate was done with 1-2 technical replicates. Technical replicates are independent pull-down assays, which used RIC-3 from the same batch of 70-110 *Xenopus laevis* oocytes. **E,** Western blot image of NACHO protein extracted from *Xenopus laevis* oocytes not injected (no-inj) and injected (NACHO) with NACHO IVT mRNA. In lanes containing NACHO injected oocyte sample (different amounts of proteins were loaded), a prominent band just below 20 kDa is detected. **F,** Western blot image of NACHO extracted from *Xenopus laevis* oocytes and NACHO in eluates from pull-down assays using 50 mM L-cysteine-capped resin and α7 or α7-A peptide covalently-linked resin. The eluates from peptide-linked resins give rise to stronger NACHO signals as compared to the eluate from the capped resin. Peptide/resin ratios in pull-down assays are ∼ 9.2 mg/ml and 5.1 mg/ml for α7 lanes (from left to right) and 6.4 mg/ml for α7-A. Data shown here are representatives of two biological replicates and at least 3 technical replicates. Biological replicates are pull-down assays conducted using different NACHO extracts from various batches, each containing 30-70 *Xenopus laevis* oocytes (two different NACHO extracts from two batches of oocytes). Each biological replicate was done with 1-2 technical replicates. Technical replicates are independent pull-down assays, which used NACHO from the same batch of 30-70 *Xenopus laevis* oocytes.

Our results indicate synthetic α7 L1-MX peptide interacts with both RIC-3 and NACHO. To the best of our knowledge this is the first time NACHO interaction has been localized to a specific domain, in this case a short segment of the ICD of α7 nAChR. When conserved sites, W330, R332, and L336 were replaced by Ala, we observed that triple and double Ala replacements abolished the biogenesis of α7nAChR, and the RIC-3 effect on the biogenesis could not be detected. In contrast, both RIC-3 and NACHO effects on the biogenesis could still be seen with the single Ala replacement, W330A, while W330A was also sufficient to abolish the functional surface expression. In addition, we detected a stable α7 pentamer in co-expression with NACHO. This NACHO-promoted α7 pentamer is sensitive to low heat but resistant to denaturing conditions and Endo H enzymes. NACHO worked synergistically with residues W330, R332, and L336 to stabilize such α7 pentamer via non-covalent interactions.

## MATERIALS AND METHODS

### L1-MX peptides

L1-MX peptides, α7, α7-AAA, and 5HT3A, Fig. 2 B, were designed based on the L1-MX segments of the α7 nACh subunit (PDB ID 7KOO and UniProt: P49582) and the 5-HT_3A_ subunit in our previous study (Do and Jansen, 2023), respectively. These L1-MX peptides were engineered to include an N-terminal cysteine (Cys) with a free sulfhydryl (−SH) group, enabling the formation of a stable thioether linkage with the iodoacetyl group of the resin (#53155; Thermo Fisher Scientific). Any naturally occurring cysteine residues were replaced with serine to prevent unwanted disulfide bond formation.

L1-MX peptides and alanine-substituted L1-MX peptides (Fig. 2 B) were synthesized and validated by GenScript and/or Biomatik. α7 and α7-AAA peptides were dissolved in DMSO, and 5HT3A peptide was dissolved in coupling buffer (50 mM Tris and 5 mM EDTA-Na, pH 8.5). To prevent oxidation of sulfhydryl groups, the reducing agent tris (2-carboxyethyl) phosphine (TCEP) was added to solubilized peptides at a final concentration of 2-2.5 mM.

After dissolution, the concentration of each peptide was determined using the following equation:

Peptide concentration (mg/ml) = (absorbance/extinction coefficient) × peptide molecular weight.

Absorbance at 280 or 205 nm was measured using a Nanodrop device (Nanodrop OneC, Thermo Fisher Scientific). The peptide extinction coefficients and molecular weights were calculated using the Expasy ProtParam webserver or following a previous study (Anthis and Clore, 2013).

### Coupling of L1-MX peptides with iodoacetyl resin

The detailed procedure for coupling L1-MX peptides with iodoacetyl resin is described in our previous study and the manufacturer’s manual (Do and Jansen, 2023). Briefly, 30 μl of UltraLink iodoacetyl resin (#53155; Thermo Fisher Scientific), settled in a Pierce spin column (#69705; Thermo Fisher Scientific), was washed three times with 300 μl of coupling buffer (50 mM Tris and 5 mM EDTA-Na, pH 8.5). Each peptide solution was then added to the resin to achieve a ratio of resin-bound peptide/resin between 5.9 mg/ml – 9.2 mg/ml. To prevent nonspecific binding between RIC-3 and the resin, the remaining active binding groups of the iodoacetyl resin were blocked using a freshly prepared solution of 50 mM L-cysteine containing 25 mM TCEP at pH 8.5. The resin was then processed according to the manufacturer’s instructions.

### Expression of RIC-3 and NACHO proteins in *Xenopus laevis* oocytes for extracting total membrane proteins

The pGEMHE19 plasmid containing the human RIC-3 gene (GenBank accession no. NM_001206671.2) was a gift from Dr. Millet Treinin (Hebrew University, Jerusalem, Israel). The plasmid vector pXOON-TMEM35 was generated by inserting the TMEM35 gene (GenBank: BC050273.1), which encodes the NACHO protein, into the pXOON vector using the BamHI and EcoRV restriction sites. To generate linear DNA template for in-vitro transcription into messenger RNA (IVT mRNA), 30 µg of plasmid vector was linearized using NheI restriction enzyme (Cat #R0131L, New England Biolabs) and CutSmart buffer (Cat #B7203S, New England Biolabs) as previously described (Do and Jansen, 2023). IVT mRNAs were synthesized using the mMESSAGE mMACHINE T7 kit (Cat # AM1344 and AM1340, Invitrogen by Thermo Fisher Scientific) and subsequently purified using the MEGAclear kit (Cat # AM1908, Invitrogen by Thermo Fisher Scientific). The resulting IVT mRNAs were dissolved in nuclease-free water and stored at -80 °C until use.

Defolliculated *Xenopus laevis* oocytes were purchased from Ecocyte Bioscience US LLC. Each oocyte was injected with 50.6 nl of RIC-3 (10 ng) or NACHO (5 ng) IVT mRNAs under a light microscope using an MM-3 micromanipulator (Narishige), a NANOJECT II device (Drummond Scientific Company), and injection glass pipettes (Cat # 3-000-203-G/X, Drummond Scientific Company) prepared with a P-97 micropipette puller (Sutter Instrument Co). The microinjection was performed in OR2 buffer (82.5 mM NaCl, 2 mM KCl, 1 mM MgCl2, 5 mM HEPES, pH 7.5). The oocytes were kept in the OR2 buffer for 15 minutes before being transferred to standard oocyte saline (SOS) medium (100 mM NaCl, 2 mM KCl, 1 mM MgCl_2_, 1.8 mM CaCl_2_, and 5 mM HEPES; pH 7.5), supplemented with 5% horse serum (Cat. # 26050088, Sigma-Aldrich) and 1 × antibiotic-antimycotic (Cat. # 15240-062, ThermoFisher Scientific/Gibco), and maintained at 15°C. The SOS medium was replaced 12 hours post-injection and then every 24 hours until the oocytes were used for experiments. Three (for RIC-3) or five (for NACHO) days after injection, oocytes were harvested for the extraction of total membrane proteins containing RIC-3 or NACHO.

### Isolation and solubilization of *Xenopus laevis* oocyte membranes containing RIC-3 or NACHO

The detailed procedure for isolating and extracting total membrane proteins containing RIC-3 or NACHO protein was described in our previous study (Do and Jansen, 2023). Briefly, *Xenopus laevis* oocytes were homogenized using a glass-Teflon homogenizer in lysis buffer (25 mM MOPS, 1 mM EDTA, and 0.02% NaN3; pH 7.5) supplemented with protease inhibitor cocktail II (SKU # P50800-1, Research Products International Corp). Cell debris was removed by centrifugation at 1,000 × *g* for 10 min at 4 °C, and the supernatant was collected and centrifuged again at 200,000 × *g* for 45 min at 4 °C to collect the membrane pellet. The pellet was then washed in lysis buffer containing 1 M NaCl and 2 mM MgCl_2_ and centrifuged again at 200,000 × *g* for 45 min at 4 °C. Afterward, total membrane proteins containing RIC-3 or NACHO were solubilized in solubilization buffer (1.5 % Triton X-100, 100 mM K-acetate, 40 mM KCl, 1 mM EDTA, 10 mM MOPS, 0.02 % NaN3, and 2 mM NEM; pH 7.5) supplemented with protease inhibitor cocktail. Solubilization was carried out at 4 °C with gently shaking. The sample was then centrifuged at 30,000 × *g* for 45 minutes at 4 °C, and the resulting supernatant was rapidly frozen in liquid nitrogen and stored at −80 °C until use.

### Pull-down assay

The detailed procedure for pull-down assays of L1-MX peptides and RIC-3 or NACHO was described in our previous study (Do and Jansen, 2023). Briefly, the resin was washed three times with 300 μl of binding buffer (0.5 % Triton X-100, 100 mM K-acetate, 40 mM KCl, 1 mM EDTA, 10 mM MOPS, and 0.02 % NaN3; pH 7.5). RIC-3 or NACHO solubilized in 1.5 % Triton X-100 as described above was thawed on ice and centrifuged at 30,000 × g for 1 hour at 4 °C before the incubation of the RIC-3-containing or NACHO-containing extract with the resin. The mixture was incubated at 4 °C for 2 hours. To remove nonspecifically bound RIC-3 or NACHO and other unbound proteins, the resin was washed with 150 μl of washing buffer 1 (1.2 % Triton X-100, 100 mM K-acetate, 40 mM KCl, 1 mM EDTA, 10 mM MOPS, and 0.02 % NaN_3_; pH 7.5), followed by an additional wash with 60 μl of washing buffer 2 (1 M NaCl, 25 mM Tris-Cl, and 1 mM EDTA; pH 7.5).

RIC-3 or NACHO protein was then eluted from the resin-bound peptides by applying 40 μl of elution buffer (277.8 mM Tris-HCl, pH 6.8, 44.4 % glycerol, 10 % SDS, 10 %

2-mercaptoethanol, and 0.02 % Bromophenol Blue) and incubating at 75°C for 5 min, followed by centrifugation at 8,000× *g* for 3 min at RT. The flow-through was analyzed using western blot/immunoblot, as described below, to detect RIC-3 or NACHO.

### Mutagenesis

The human α7 AChR subunit (CHRNA7, GenBank accession no. NM_000746.6) in the plasmid vector pMXT (MX-wt) was a gift from Dr. Jon M. Lindstrom (University of Pennsylvania, Philadelphia, PA). Using this plasmid as a template, triple (W330A, R332A, and L336A), double (R332A and L336A), and single (W330A) amino acid substitutions in the L1-MX segment were introduced using the QuikChange II XL Site-Directed Mutagenesis Kit (Cat # 200522 Agilent Technologies) and oligonucleotide primers (Sigma). The DNA sequences containing these substitutions, MX-AAA, MX-AA, and MX-A, were verified by sequencing (Genewiz/Azenta). Amino acid numbering follows UniProt P36544.

### Expression of *α*7 nACh channels in the presence and absence of NACHO and RIC-3 in *Xenopus laevis* oocytes for TEVC recordings and cell surface labeling

Four plasmid constructs, MX-wt, MX-AAA, MX-AA, and MX-A, were linearized using BamHI restriction enzyme (Cat # R0136L, New England Biolabs) and NEB 3.1 buffer (Cat # B7203S, New England Biolabs). Plasmid vectors pXOON-TMEM35 and the pGEMHE19 plasmid containing the human RIC-3 gene were linearized using NheI restriction enzyme and CutSmart buffer. The linearized DNA plasmids were then used to produce IVT mRNA, as described for RIC-3 above.

Ten nanograms of MX-wt, MX-AAA, MX-AA or MX-A IVT mRNA were injected into a *Xenopus laevis* oocyte in the presence or absence of 5 ng of RIC-3 IVT mRNA and/or 5 ng of NACHO IVT mRNA. The injection of IVT mRNAs and the subsequent maintenance of oocytes post-injection are described above. Five days post-injection, the oocytes were used for two-electrode voltage-clamp (TEVC) recordings or cell surface labeling.

### Electrophysiology – *Xenopus laevis* oocyte interaction assay

Changes in the functional surface expression of α7 nACh channels in the presence and absence of chaperone proteins RIC-3 and/or NACHO were probed using the two-electrode voltage-clamp (TEVC) recordings 5 days post-injection of oocytes.

Each oocyte was placed in a 200 μl perfusion chamber containing the recording buffer (82.5 mM NaCl, 1 mM MgCl_2_, 20 mM KCl, and 5 mM HEPES, 2 mM CaCl2; pH 7.5), which was perfused into the chamber at a rate of 2-3 ml/min. The ground electrode, containing 1% agarose in 3 M KCl, was connected to the bath. Voltage and current electrodes were filled with 3 M KCl solution, with resistances ranging from 1 to 5 MΩ. Oocytes were voltage-clamped at a holding potential of -60 mV using a TEV-200A Voltage Clamp amplifier (Dagan Corporation). Data were acquired using a Digidata 1440A digitizer (Molecular Devices) and Clampex 10.7 software in gap-free acquisition mode and with a 200 Hz sampling rate. Currents of α7 nACh channels were evoked by applying 1 mM acetylcholine chloride (Cat # 159171000, Thermo Scientific Chemicals) in recording buffer.

### Cell surface labeling

Biotinylation of extracellular amines of membrane proteins was carried out as described in our previously published studies (Jansen et al., 2008; Duddempudi et al., 2013; Date et al., 2016). Oocytes expressing MX-wt, MX-AAA, MX-AA, or MX-A α7 nACh channels, either alone or co-expressed with RIC-3 or NACHO, as described above, were washed with CFFR buffer (115 mM NaCl, 1.8 mM MgCl2 x 6H2O, 2.5 mM KCl, 10 mM HEPES; pH 8.0) to remove SOS medium, horse serum, and antibiotic-antimycotic. While the oocytes were kept in CFFR buffer, EZ-Link Sulfo-NHS-LC-Biotin (Prod # 21335, Thermo Scientific) was added at a final concentration of 0.5 mg/ml. The biotinylation was carried out for 30 minutes at room temperature. Excess biotinylating agent was removed by thoroughly washing the oocytes with Buffer H (20 mM Tris, 100 mM NaCl; pH 7.4).

Next, the same number of oocytes from each expression condition was transferred to a 1.5 ml microcentrifuge tube carrying over a minimal amount of buffer solution. The oocytes were then homogenized in ice-cold Buffer H+++ at 40 µl / oocyte (Buffer H++: 20 mM Tris, 100 mM NaCl, 1% Triton, 0.5% sodium deoxycholate, 10 mM NEM, pH 7.4, supplemented with protease inhibitor cocktail II (SKU # P50800-1, Research Products International Corp) at a ratio of 20 µl cocktail per 1 ml of Buffer H++ by repeated pipetting with a 1 ml pipette tip until no particles or dark granules remained. Subsequently, cell lysis and protein extraction were carried out for 1 hour at 4 °C by gentle mixing with inversion. To separate cell debris, the samples were then centrifuged at 20,000 × *g* for 10 min at 4 °C. The middle layer (supernatant) between the floating white lipid layer and pelleted yolk was moved into a new ice-cold 1.5 mL microcentrifuge tube. The pellet was resuspended in the ice-cold Buffer H+++ at 20 µl / oocyte, and the centrifugation was repeated. The second supernatant was combined with the first. More ice-cold Buffer H+++ was then added to this solution to obtain the total proteins of each condition at 70 µl /oocyte.

Biotinylated proteins were then isolated from the remaining total protein samples using pre-washed NeutrAvidin beads (High capacity neutravidin agarose resin, #29204, Thermo Scientific product). Prior to use, 75 µl of settled NeutrAvidin beads in a 1.5 ml microcentrifuge tube were washed three times at room temperature with 1 ml of Buffer H+ (20 mM Tris, 100 mM NaCl, 1% Triton, 0.5% sodium deoxycholate; pH 7.4). After each wash, the beads were pelleted by centrifugation at 2,500 x g for 2 min, and the wash solution was then removed.

The mixture of total proteins and NeutrAvidin beads was incubated at 4 °C with gentle shaking for 90 min. After incubation, the beads were separated from the supernatant by centrifugation at room temperature at 2,500 x g for 2 min. The supernatant was carefully removed. Next, the beads were washed three times: the first two washes were performed with 1 ml of Buffer H+ and rotating samples at room temperature for 10 min. After each wash, the supernatant was removed. In the final wash, Buffer H+ containing 2 % SDS was used instead of Buffer H+ alone. Biotinylated proteins were eluted from the beads by adding 4x Laemmli buffer (277.8 mM Tris-HCl, pH 6.8, 44.4 % glycerol, and 0.02 % Bromophenol Blue) containing either 8 % SDS and 10% 2-mercaptoethanol or 1x Laemmli buffer containing 1% SDS and 1 mM DTT, at ∼3 µl /oocyte and incubating at 37 °C for 10 min. The eluted proteins were separated from the beads after centrifugation at 2,500 x g for 2 min at room temperature. The equal volume from each sample was used for immunoblot analysis.

### Immunoblot analysis

The detailed protocol for the detection of RIC-3 using a primary/mouse antibody (Cat. #H00079608-B01P; Abnova Corporation) and a goat-raised anti-mouse secondary antibody (#31430; Thermo Fisher Scientific) was described in our previous study (Do and Jansen, 2023). Briefly, the eluates from the pull-down assay were separated using sodium dodecyl sulfate–polyacrylamide gel electrophoresis (SDS-PAGE). The SDS-PAGE was conducted at a constant voltage of 180 for 35 - 40 minutes at room temperature using precast gels (Cat #4561094, #4561096, or #4561023 Bio-Rad) and running buffer Tris/Glycine/SDS (Cat #1610772, Bio-Rad). The proteins were then transferred to polyvinylidene fluoride (PVDF) membranes (Bio-Rad) using the Trans-Blot Turbo RTA transfer kit (Cat #1704272, Bio-Rad) following the manufacturer’s protocol.

PVDF membranes were then blocked under agitation in 5 % non-fat milk in Tween-containing Tris-buffered saline buffer (TTBS buffer: 0.1 % Tween-20, 100 mM Tris, and 0.9 % NaCl, pH 7.5) for 1 hour at room temperature or overnight at 4 °C. The blot was then incubated with the RIC-3 primary antibody in a dilution of 1:2,000, overnight at 4 °C with gentle shaking. Each blot was then washed in TTBS buffer before the incubation with goat-raised anti-mouse secondary antibody, conjugated to horseradish peroxidase enzyme, HRP (#31430; Thermo Fisher Scientific product), in a dilution of 1:5,000 for 2 hours at room temperature, under gentle agitation.

The protocol for detecting NACHO protein is similar to the procedure described above for RIC-3, except for the antibodies. We used anti-NACHO primary/rabbit raised antibody (ThermoFisher, Cat # PA5-61774) at a dilution of 1:1,000 dilution and goat-raised anti rabbit secondary antibody conjugated to Horseradish peroxidase enzyme (ThermoFisher Cat # 31460, at a dilution of 1:20,000 dilution for 1.5 hours at room temperature.

To detect α7 nACh oligomers, the biotinylated proteins from the biotinylation assays were also separated using the SDS-PAGE method using Mini-PROTEAN TGX Gels (Cat #456-1094 and #4561023, BioRad products) as described above. The proteins were then transferred from the gels to PVDF membranes. The PVDF blots were then blocked under agitation in the TTBS buffer containing 5 % non-fat milk at 4 °C overnight. Next, the blots were incubated with the rat-raised anti α7 nACh primary antibody (Cat # sc-58607, 200 µg/ml, Santa Cruz product) in a dilution of 1:400 with TTBS buffer containing 5% non-fat milk for 20-22 h at 4°C with gentle agitation. Each blot was then washed three times with TTBS buffer, each for 5 minutes, at room temperature and 80 or 90 rpm shaking. After the washes, the blots were incubated with goat-raised anti-rat secondary antibody conjugated to horseradish peroxidase enzyme (# ab97057, Abcam product) in a dilution of 1:5,000 or 1:20,000 dilution with TTBS buffer containing 5% non-fat milk for 1.5 hours, 70 rpm shaking at room temperature.

After the incubation with secondary antibodies, each blot was washed 5 times with TTBS buffer and once with tris-buffered saline (100 mM Tris and 0.9 % NaCl, pH 7.5). The chemiluminescent detection for RIC-3 and α7 nACh proteins was done using SuperSignal West Femto Maximum Sensitivity Substrate (# 34095, Thermo Fisher Scientific product) and a digital imaging system (ChemiDoc MP, Bio-Rad).

### Cross-linking assay with CuSO_4_ and Phenanthroline

Phenanthroline (Sigma Aldrich, Cat # 13137-25G) was prepared as a 1 M stock solution in DMSO and CuSO_4_ (Sigma Aldrich, Cat # 203165-25G) as a 100 mM stock solution in water. CuSO_4_ and Phenanthroline were mixed in deionized water) before use to achieve working concentrations of 10 mM : 20 mM Cu:Phen.

After incubating the oocytes that expressed the MX-wt construct alone with EZ-Link Sulfo-NHS-LC-Biotin (Prod # 21335, Thermo Scientific), the oocytes were thoroughly washed with Buffer H. These oocytes were then incubated with Cu:Phen at final concentrations of 10 µM and 20 µM, respectively, for 8 minutes at room temperature. Afterward, the Cu:Phen was washed out thoroughly with Buffer H, and the extraction of the total protein was continued as described in the cell surface labeling section.

### Statistical Analysis

For functional surface expression or TEVC data, α7 current amplitude was obtained by subtracting the peak current (achieved after applying of 1 mM acetylcholine) from the resting current (before the application of 1 mM acetylcholine). The current amplitudes from different experimental conditions were then normalized with the current amplitude obtained from the condition of MX-wt + NACHO of the same batch of *Xenopus* oocytes. For surface expression of α7 pentamers, relative quantification of α7 pentamers was done using Image Lab (Bio-Rad) application. Each blot was analyzed to identify the lanes and bands of interest as described in the user’s manual. The peak volume of each band was normalized to the peak volume of α7 pentameric band from MX-wt + NACHO sample from the same batch of *Xenopus* oocytes. We used column as the data table format in GraphPad Prism 10.6.1 to enter normalized data previously done in Microsoft Excel 2021. Data from an experimental condition were entered with replicate values and stacked into a column. The statistical significance was determined using one-way ANOVA with Tukey’s or Dunnett’s multiple comparisons test integrated in the GraphPad Prism 10. The standard error of the mean was calculated and reported as the error bar.

## RESULTS

We have recently shown that RIC-3 affects the functional surface expression of 5-HT_3A_ channels via the motif D**W**L**R**…V**L**DR. This motif was identified in the L1-MX segment and the MA-M4 junction of the 5-HT_3A_-ICD (Jansen et al., 2008; Pirayesh et al., 2020; Do and Jansen, 2023). Comparing the L1-MX and MA-M4 segments of the 5-HT_3A_ subunit with those of the α7 nACh subunit reveals that Trp, Arg, and Leu residues in the RIC-3 motif are conserved in L1-MX (Fig. 2 A). However, unlike the 5-HT_3A_ subunit, the α7 subunit does not contain identical conserved residues in its MA-M4 junction. The similarities and differences in the existence of the conserved residues in L1-MX and MA-M4 segments raise questions about the roles of conserved Trp, Arg, and Leu residues in the assembly and functional surface expression of α7 nAChR. Additionally, previous studies have reported that RIC-3 chaperone affects the functional surface expression of both 5-HT_3A_ and nACh channels (Miller et al., 1996; Halevi et al., 2002; Castillo et al., 2005; Cheng et al., 2005; Lansdell et al., 2005; Williams et al., 2005; Jansen et al., 2008; Wang et al., 2009; Walstab et al., 2010; Dau et al., 2013), prompting us to investigate the involvement of these conserved residues in the communication between α7 subunits and chaperone proteins such as RIC-3 and NACHO, which enhance the biogenesis of α7 nAChR.

### Synthetic *α*7 nACh L1-MX peptide interacts with RIC-3 and NACHO

To examine the relationship between the α7 subunit and RIC-3 and NACHO, we employed a pull-down assay using a synthetic α7 L1-MX peptide and RIC-3- or NACHO-containing extracts. The α7 L1-MX peptide (Fig. 2 B, α7), with an added N-terminal Cys, was covalently linked to iodoacetyl resin. First, the peptide-linked resin was used to capture RIC-3 from protein extracts of *X. laevis* oocytes injected with RIC-3 IVT mRNA. RIC-3 was detected in the protein extracts from oocytes injected with RIC-3 IVT mRNA, which is near 50 kD, but not in uninjected oocytes (Fig. 2 C). We used the synthetic 5-HT_3A_ L1-MX peptide (Fig. 2 B, 5HT3A) from our previous study as a positive control to confirm an interaction between the peptide and RIC-3 (Do and Jansen, 2023). RIC-3 signals, obtained after western blot with RIC-3 antibody, from pull-down samples using 5HT3A and α7 peptide-linked resins are stronger than RIC-3 signals from capped resins without peptides, which we infer to represent unspecific binding (Fig. 2 D). These data indicate that RIC-3 interacts with the α7 synthetic L1-MX peptide. Next, we used the α7 L1-MX peptide with Ala replacements at the conserved residues (Fig. 2 B, α7-AAA) in our pull-down assays. Similar to wild-type peptide, the RIC-3 signal from pull-down assays using α7-AAA peptide-linked resin is much stronger than the RIC-3 signal from resin alone (Fig. 2 D).

Previous findings show that RIC-3 and NACHO positively regulate the functional surface expression of α7 nACh channels (Halevi et al., 2002; Halevi et al., 2003; Castillo et al., 2005; Lansdell et al., 2005; Williams et al., 2005; Gu et al., 2016). We, therefore, examined the interaction between NACHO and α7 L1-MX peptide. The pull-downs with α7 L1-MX peptide and α7-AAA peptide were conducted with NACHO extracts obtained by injecting NACHO IVT mRNA into *X. laevis* oocytes. Fig. 2 E represents a western blot of extracts from *X. laevis* oocytes. Using a NACHO primary antibody, we observed a band near 18 kD from *X. laevis* oocytes that were injected with NACHO IVT mRNA. This 18 kDa band corresponds to the expected molecular weight of NACHO protein. However, we did not observe a similar band in uninjected oocytes. The NACHO band was also detected in eluates from pull-down assays using α7 or α7-AAA peptide-linked resins and the NACHO-containing extracts (Fig. 2 F). NACHO signals from pull-down samples with the peptide-linked resins are much stronger than NACHO signals from capped resins without peptides.

Together, our data here indicate that the α7 L1-MX peptides pull down both RIC-3 and NACHO.

### Ala substitution of conserved residues in L1-MX disrupts *α*7 nAChR’s biogenesis, and the co-expression with NACHO and/or RIC-3 partially rescues functional expression

A closer look into the 3D structure of α7 nAChR (Fig. 3 A) suggests that the side chain of residue W330A points in an opposite direction as compared to those of residue R332 and L336. Based on this observation, we generated a single Ala substitution (MX-A: W330A) and a double Ala substitution (MX-AA: R332A and L336A) in addition to a triple Ala substitution (MX-AAA: W330A, R332A, and L336A) as full-length α7 nACh subunits (Fig. 3 B).

**Figure 3:**
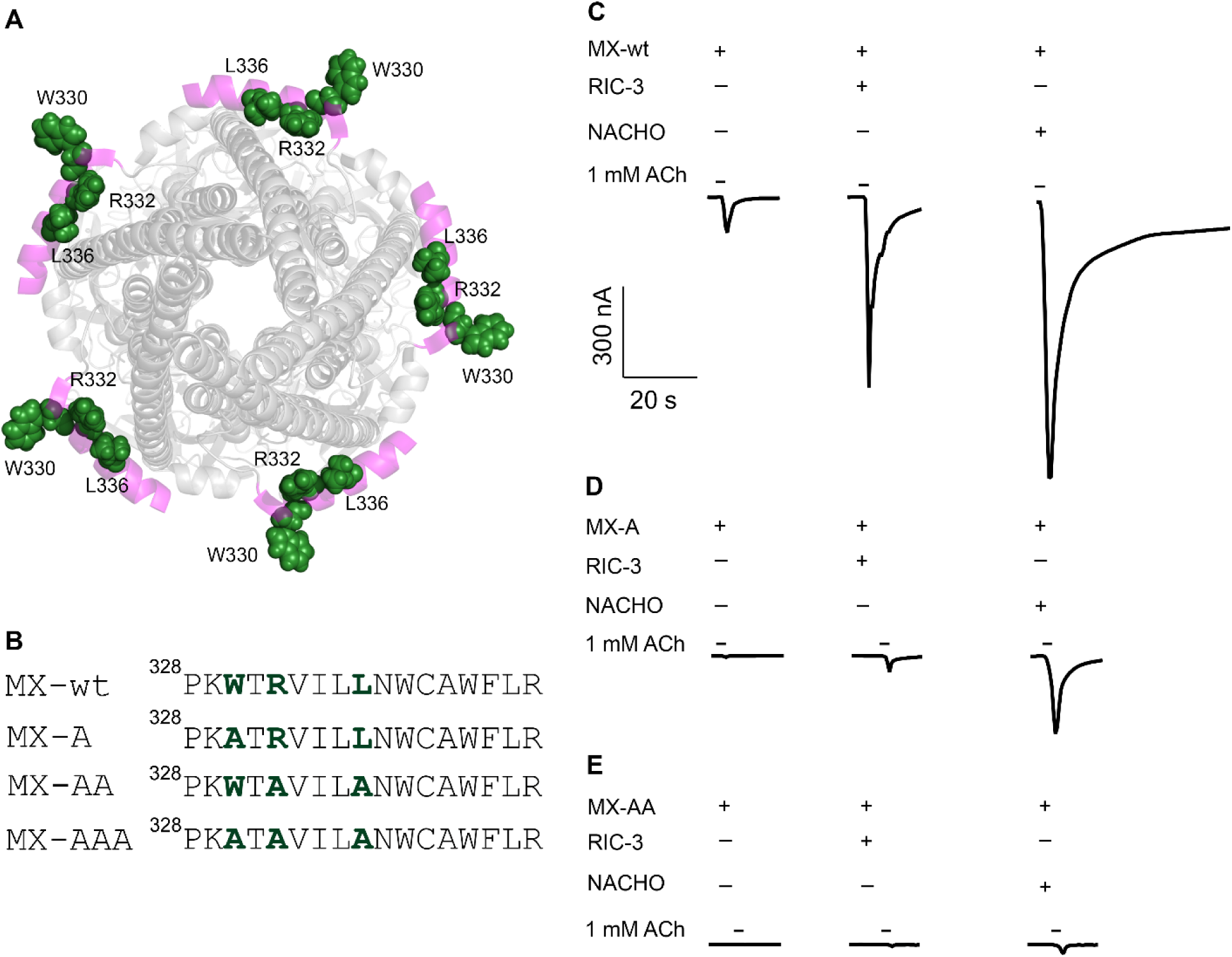
Representative acetylcholine-elicited current traces of *α*7 nACh channels alone or in the presence (+) of RIC-3 or NACHO. A, View of pentameric α7 nACh channel (PDB ID 7KOO) from the intracellular side with residues W330, R332, and L336 in green. **B,** The amino acid sequence of the MX segments in the MX-wt, MX-A, and MX-AA constructs. The residue numbers are cited as in UniProt P49582. **C, D, and E,** Representative current traces of α7 nACh channels (MX-wt, MX-A, and MX-AA constructs) alone or in the presence (+) or absence (-) of RIC-3 or NACHO.

For α7 nAChR wildtype, co-expression with RIC-3 increased α7 current amplitudes about 6-fold (Fig. 3 C and Fig. 4 A, MX-wt vs. MX-wt + RIC-3, one-way ANOVA with Tukey’s multiple comparisons test, P < 10^-10^). NACHO co-expression similarly enhanced the α7 current amplitude approximately 12-fold (Fig. 3 C and Fig. 4A, MX-wt vs. MX-wt + NACHO, P < 10^-10^). The NACHO effect was significantly larger than that of RIC-3 (Fig. 3 C & Fig. 4 A, MX-wt + RIC-3 vs. MX-wt + NACHO, P < 6 x 10^-14^). We did not observe synergism between NACHO and RIC-3 (Fig. 4A, MX-wt + NACHO vs. MX-wt + RIC-3 + NACHO) as previously shown in HEK293T cells (Gu et al., 2016; Dawe et al., 2019).

**Figure 4.**
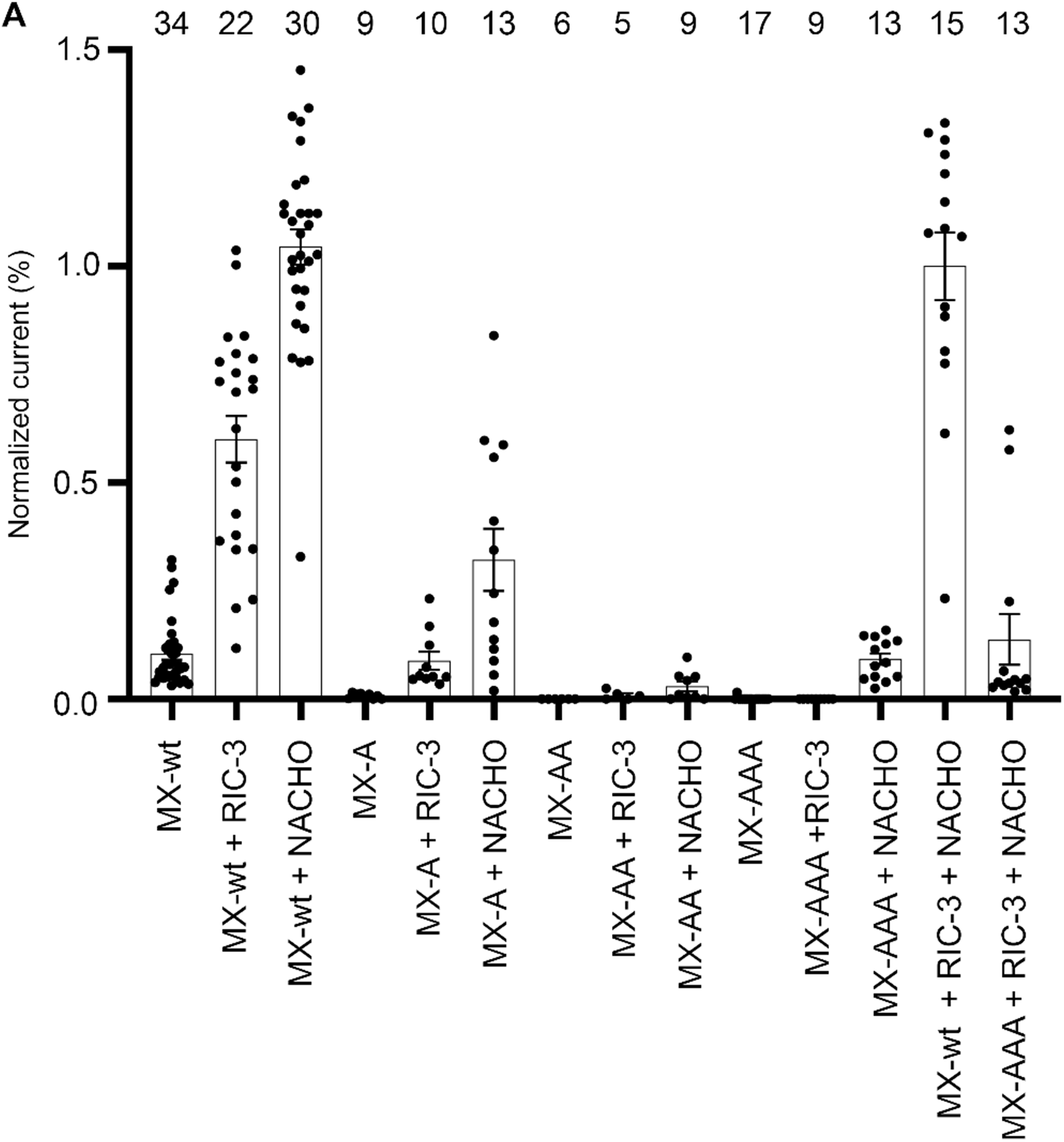
Ala substitutions at residues W330, R332, and L336 in the MX segment of *α*7 nACh subunit disrupt the functional surface expression of *α*7 nAChR. 1 mM acetylcholine-induced normalized currents elicited from MX-wt, MX-A, MX-AA and MX-AAA constructs expressed alone in *Xenopus laevis* oocytes or co-expressed with RIC-3 and/or NACHO. Biological replicates are numbers at the top of the chart. Statistical significance was determined using ordinary one-way ANOVA with Tukey’s multiple comparisons test. Statistical significance for MX-wt vs. MX-wt + RIC-3, MX-wt vs. MX-wt + NACHO, and MX-wt + RIC-3 vs. MX-wt + NACHO with P = 3.9 x 10^-14^, P < 10^-10^, and P < 10^-10^, respectively. There was no significant difference in α7 current amplitude when co-expressed with NACHO or both NACHO and RIC-3. The α7 current amplitude elicited from the MX-wt + RIC-3 and MX-wt + NACHO conditions significantly differ from those of the MX-A + RIC-3 and MX-A + NACHO conditions, respectively (P= 0.4 x 10^-10^, P < 10^-11^).

No currents were observed for the MX-A, MX-AA, and MX-AAA constructs in the absence of RIC-3 or NACHO (Fig. 3 D and E, Fig. 4 A), indicating that these substitutions completely disrupt biogenesis of functional channels.

However, for MX-A co-expressed with RIC-3 or NACHO, we did observe currents. Compared to the wildtype co-expressed with NACHO or RIC-3, the current amplitude elicited for the MX-A construct was reduced by ∼72% and ∼51%, respectively (Fig. 4 A, MX-wt + NACHO vs. MX-A + NACHO, P < 10^-11^; MX-wt + RIC-3 vs. MX-A + RIC-3, P = 4 x 10^-11^). Since there was no current with MX-A alone, it is not possible to calculate the size effect of the current increase. However, clearly, either chaperone led to a rescue of MX-A functional expression.

Similarly, we observed very small currents for the MX-AA construct after co-expression with RIC-3 or NACHO. We did not detect any current for the MX-AAA construct after co-expression with RIC-3, however, some current was detected after co-expression with NACHO (Fig. 4 A).

Our findings suggest that Ala replacements of conserved residues in L1-MX impair the functional biogenesis of α7 channels, while NACHO and/or RIC-3 co-expression leads to a rescue of functional expression

### NACHO stabilizes *α*7 pentamer via non-covalent interactions

Five days post-expression in *Xenopus* oocytes, cell-surface α7 proteins were labeled with biotin and isolated using NeutraAvidin beads (see the Method section for more details). After SDS PAGE separation and Western blot, a prominent band containing biotinylated α7 proteins appeared at approximately 250 kDa on a 7.5% polyacrylamide gel (Fig. 5 A). The predicted monomer size of the α7 protein is 56 kDa. We infer the prominent band to represent homopentameric assemblies. We observed that NACHO significantly enhanced the cell-surface expression of α7 pentamer as compared with RIC-3 (Fig. 5A & B, MX-wt + RIC-3 vs. MX-wt + NACHO, ordinary one-way ANOVA with Dunnett’s multiple comparisons test, P < 6.1 x 10^-4^). This result aligns with our TEVC data shown above (Fig. 4). However, in contrast with our TEVC data, where MX-wt alone produces lower α7 current amplitude compared with MX-wt co-expressed with RIC-3 or NACHO, MX-wt alone construct gives rise to the highest level of the cell-surface expression of the α7 pentamer compared to MX-wt with RIC-3 or NACHO (Fig. 5A & B, MX-wt vs. MX-wt + RIC-3 or MX-wt + NACHO). In addition, the intensity of this pentameric band is significantly decreased in MX-A and MX-AA constructs compared with that from the wild-type MX-wt construct (Fig. 5 A & B, MX-A + NACHO / MX-AA + NACHO vs. MX-wt + NACHO or MX-wt, ordinary one-way ANOVA with Dunnett’s multiple comparisons test).

**Figure 5.**
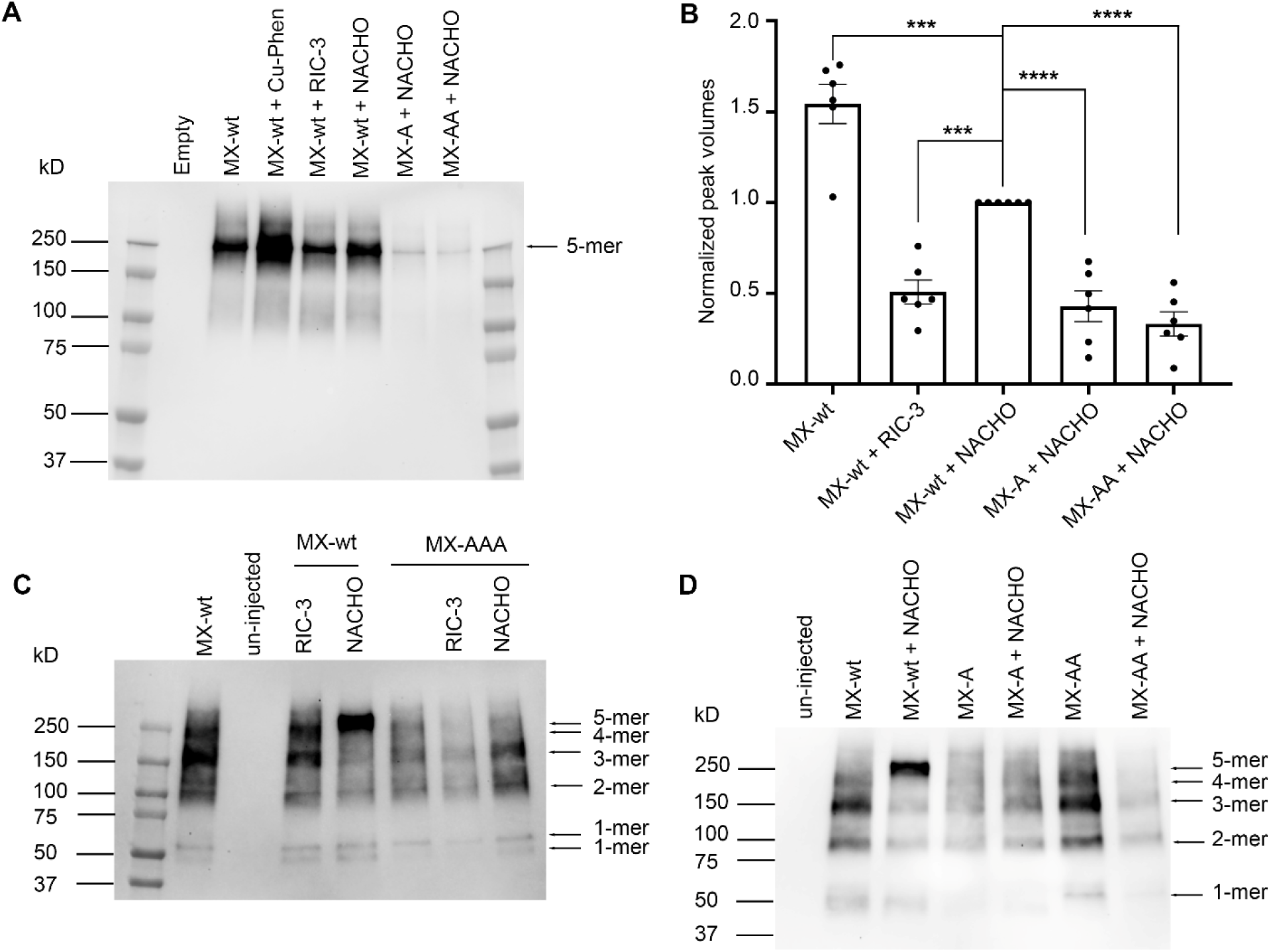
NACHO and L1-MX conserved residues promote the stabilization of *α*7 pentamers. A, Cell surface biotinylated α7 oligomers from MX-wt, MX-A (W330A) and MX-AA (R332A & L336A) constructs in co-expression with and without RIC-3 / NACHO. The biotinylated α7 proteins were eluted from the NeutrAvidin beads using Laemmli buffer containing 1 % SDS and 1 mM DTT at 37°C for 10 minutes. Samples were separated on the 7.5% polyacrylamide gel. Biotinylated proteins from six oocytes were loaded in a lane. The data shown here is the representative of three biological replicates and 6 technical replicates. Each biological replicate contains data collected from a batch of *Xenopus* oocytes. Each technical replicate is a western blot. The “5-mer” indicates α7 pentamer. The “Cu:Phen” indicates the samples were treated with 10 µM CuSO_4_ (Cu) and 20 µM phenanthroline (Phen). **B,** Relative quantification of α7 pentamers as presented in panel A using Image Lab (Bio-Rad) application. The peak volume of each band was normalized to the peak volume of α7 pentameric band from MX-wt + NACHO sample. The data shown here are from three biological replicates and 6 technical replicates. Each biological replicate contains data collected from a batch of *Xenopus* oocytes. In total, three different batches of the oocytes were used. Each technical replicate is a western blot. Statistical significance was determined using ordinary one-way ANOVA with Dunnett’s multiple comparisons test. Star signs, ***, indicate a P value < 6.1 x 10^-4^. Star signs, ****, indicate a P value < 8.3 x 10^-5^. The error bars are represented the standard errors of the means. **C & D,** Cell surface biotinylated α7 oligomers isolated from MX-wt, MX-AAA ((W330A, R332A, & L336A), MX-A (W330A), and MX-AA (R332A & L336A) constructs in co-expression with RIC-3 or NACHO. The biotinylated α7 proteins were eluted from the NeutrAvidin beads using Laemmli buffer containing 8 % SDS and 10 % 2-mercaptoethanol at 37 °C for 10 minutes. Samples were separated on the 4-20 % gradient gel. Biotinylated proteins from six oocytes were loaded in a lane. 5-mer, 4-mer, 3-mer, 2-mer, and 1-mer indicate α7 pentamer, tetramer, trimer, dimer, and monomer. The data shown here is the representative of at least three technical replicates and two biological replicates which came from two different batches of *Xenopus* oocytes.

When the biotinylated α7 proteins were eluted from the NeutraAvidin beads using a stronger denaturing condition, Laemmli buffer containing 8 % SDS and 10 % β-mecapthoethanol (BME), and separated on 4-20% gradient gels, we detected multiple bands near 50 kD, 100 kD, 150 kD, and 250 kD (Fig. 5 C & D), which are similar to the predicted molecular weights of monomer (56 kD), dimer (112 kD), trimer (168 kD), tetramer (224 kD), and pentamer (280 kD) of α7 proteins (Fig. 5 C & D). Interestingly, under these conditions, we consistently observed the strongest pentameric band from MX-wt + NACHO. This band is absent or much less pronounced in other conditions (MX-wt and MX-wt + RIC-3) when treated under these denaturing conditions (Fig. 5 C & D). These data demonstrate that NACHO promotes the stabilization of the α7 pentamer.

The human α7 ICD contains seven Cys residues; however, only three were visible in the experimental structure (PDB ID 7KOO). Their sulfhydryl (SH) groups are separated by a range of 13 Å to 28 Å. The remaining four Cys residues are located in the loop connecting L1-MX and M4. Based on our observation of denaturation-resistant pentameric bands upon co-expression with NACHO, but not other expression conditions, we infer that NACHO may promote the formation of disulfide bonds among intracellular Cys residues, thereby stabilizing the α7 pentamer. If this were the case, we would be able to promote the generation of such intra- or intermolecular disulfide bonds using an oxidizing agent that catalyzes the formation of disulfide bonds between SH groups (Kobashi, 1968). Copper phenanthroline-induced oxidation to form intramolecular disulfide bonds has previously been used for structure-function studies of pLGICs (Reeves et al., 2005; Bali et al., 2009; Mnatsakanyan and Jansen, 2013). Here, we treated the α7 sample extracted from the MX-wt construct expressed alone with copper phenanthroline (Cu:Phen) as described in the Methods. We then treated both the Cu:Phen treated α7 proteins and the NACHO-promoted α7 proteins under the same denaturing conditions. We observed distinct dissociation patterns between the Cu:Phen α7 and NACHO-promoted α7 pentamers: the most prominent band in the NACHO-promoted α7 samples is the pentamer, which is not the case in the Cu:Phen α7 samples (Fig. 6 A). Our results show that oxidizing conditions alone are not sufficient to cause stable pentamers to the same degree as compared to those obtained by NACHO co-expression. It is possible that NACHO enables inter- or intramolecular disulfide bonds, however we do not have experimental evidence to support this statement.

**Figure 6.**
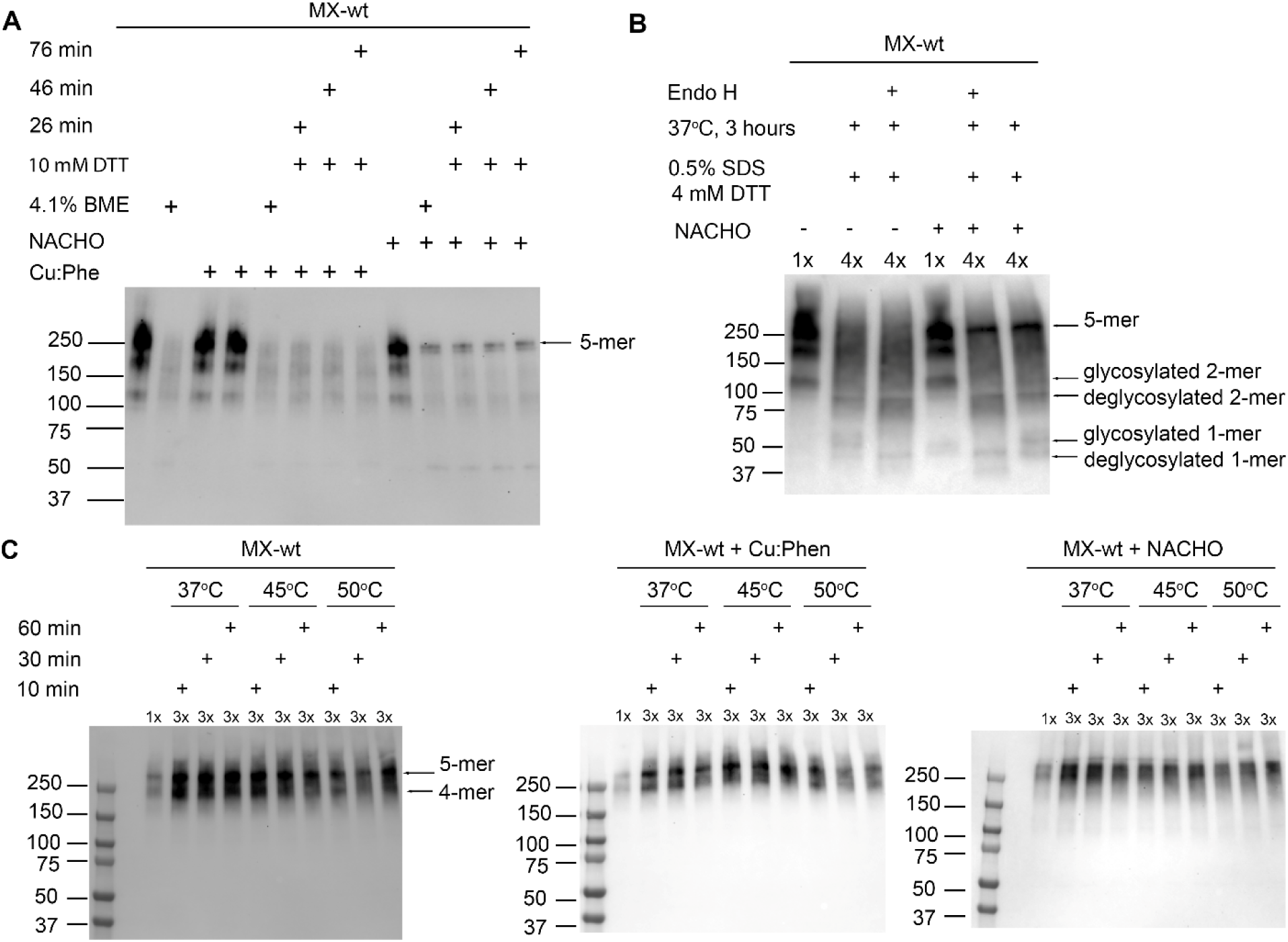
NACHO-promoted *α*7 pentamer is resistant to strong denaturing condition but sensitive to low temperatures. A, NACHO-promoted α7 pentamer (5-mer) is resistant to strong denaturing conditions of 3.3% SDS and 10 mM DTT or 4.1% 2-mercaptoethanol (BME) at different time points (26, 46, and 76 minutes). Cu:Phen indicates the samples were treated with 10 µM CuSO_4_ (Cu) and 20 µM phenanthroline (Phe). 10 % (v/v) of the total protein extract from an oocyte was loaded for every sample. **B,** NACHO-promoted α7 pentamer (5-mer) is resistant to the digestion of Endo H enzyme. After the Endo H treatment, the total protein samples were mixed with 4x Laemmli buffer containing 4% SDS and 4 mM DTT before SDS-PAGE and western blot. **C,** NACHO-promoted α7 oligomers were prone to aggregation at low temperatures. Left panel and middle panel show two distinctive tetrameric and pentameric band, which isn’t the case for the right panel, at which, the treatment with low heat cause tetramers and pentamers aggregates. 1x and 3x indicate 2 µg and 6 µg of total proteins were used. Total protein concentration was determined by microBCA assay. The samples were mixed with SDS-PAGE sample buffer with the final concentrations of 1% SDS and 1 mM DTT. All samples separated on 4-20% gradient polyacrylamide gels before blotting for the antibody detections. 5-mer, 4-mer, 3-mer, 2-mer, and 1-mer are abbreviated version of pentamer, tetramer, trimer, dimer, and monomer. The plus (+) and minus (-) signs indicate the presence or absence of the treated conditions, respectively.

Previous studies demonstrate that α7 nAChR is a glycoprotein and that NACHO promotes N-glycosylation in α7 nAChR (Chen et al., 1998; Gu et al., 2016; Kweon et al., 2020). N-glycosylation is critical in stabilizing protein folding and structures (Duran-Romana et al., 2024; Whittington et al., 2024; Guay et al., 2025; Hao et al., 2025). We thus hypothesize that, the added glycans to α7 nAChR help stabilize the NACHO-promoted α7 pentamers. To test this, we treated the α7 proteins and NACHO-coexpressed α7 proteins with Endo H. Endo H cleaves N-linked oligosaccharides from glycoproteins but not complex glycans. We observed that a large portion of the NACHO-promoted α7 pentamer resists Endo H digestion, which is not the case for the α7 pentamer in the absence of NACHO co-expression (Fig. 6 B, lane 3 vs. lane 5). Our finding suggests that either complex glycans are attached to α7 receptors, or Endo H enzyme could not access cleavage sites between glycans and α7 subunit. It is also possible that non-covalent interactions between the glycans and surrounding molecules prevent Endo H to access cleavage sites.

Non-covalent interactions, such as hydrophobic interactions, salt bridges, and hydrogen bonds, are sensitive to low temperatures. To examine whether non-covalent interactions help stabilize the NACHO-promoted α7 pentamers, we incubated the α7 proteins, Cu:Phen treated α7 proteins, and α7 proteins co-expressed with NACHO at low heat. At moderate temperatures of 37 °C, 45 °C and 50 °C, we observed distinct pentameric and tetrameric bands from the α7 proteins and Cu:Phen treated α7 proteins (Fig. 6 C, right and middle panels). In contrast, the samples containing α7 proteins co-expressed with NACHO give rise to continuous smear stretches. A slight smear of α7 proteins co-expressed with NACHO may further indicate non-covalent NACHO stabilization. However, further experiments would be needed to delineate mechanistic insights. Our finding here suggests that the NACHO-promoted α7 pentamer is also stabilized by non-covalent interactions.

## DISCUSSION

The ICD of the pLGIC superfamily, especially the L1-MX subdomain, is a crucial structural element that tightly regulates the surface expression of nACh and 5-HT_3A_ receptors. In muscle nAChRs, several residues in the L1-MX subdomain of β, γ, and ε subunits, such as Pro, Lys, and Leu, are essential for surface expression (Rudell et al., 2020). A study on receptor trafficking reported that residues Phe and Leu in the L1-MX subdomain of the neuronal nAChR α4 and β2 subunits are required for cell-surface expression of α4β2 nACh receptors (Ren et al., 2005). In the 5-HT_3A_ channel receptor, our previous studies have shown that the maturation and functional surface expression of the receptor expressed in *Xenopus laevis* oocytes is modulated by the L1-MX and MA-M4 segments of the ICD of the 5-HT_3A_ subunit, with assistance from the RIC-3 chaperone (Jansen et al., 2008; Pirayesh et al., 2020; Do and Jansen, 2023). Our finding is consistent with a previous study in HEK-293 cells, in which the functional surface expression of 5-HT_3A_ is abolished by deleting 10 residues in the L1-MX subdomain (Baptista-Hon et al., 2013). In α7 nAChRs, studies across various cell types demonstrate the critical role of the ICD of the α7 subunit associated with the functional surface expression (Valor et al., 2002; Castillo et al., 2005; Castelan et al., 2007; Mukherjee et al., 2009; Dawe et al., 2019); however, the information about L1-MX roles remains incomplete.

In this work, we explored the regulatory roles of residues W330, R332, and L336 in the biogenesis of α7 nAChR in the presence and / or absence of RIC-3 and NACHO chaperones. These residues are conserved and located at the L1-MX segment of the ICD of the α7 nACh subunit. These conserved residues are also found within the RIC-3 binding motif in the 5-HT_3A_ subunit in our previous study (Do and Jansen, 2023).

Our pull-down data indicate that synthetic α7 L1-MX peptides interact with both RIC-3 and NACHO (Fig. 2 D & H). However, residues W330, R332, and L336 are not crucial for the interactions (Fig. 2 F & H). Our finding here on RIC-3 differs from our previous study, in which substitution of these conserved residues in the 5-HT_3A_ L1-MX segment abrogated interaction with RIC-3 (Do and Jansen, 2023).

Using full-length receptors, we find that the triple Ala replacement at residues W330, R332, and L336 abolishes α7 current (Fig. 4 A). By contrast, the triple Ala replacement at their counterparts W347, R349, and L353 in the 5-HT_3A_ L1-MX segment reduces the serotonin current by half (Do and Jansen, 2023). In addition, the single Ala replacement (W330A) in α7 nAChR also abolishes α7 current (Fig. 3 D and Fig. 4 A), whereas the single Ala replacement (W347A) maintain the serotonin current. Our result here indicates that the conserved sites in the α7 L1-MX segment play distinctive roles as compared to their counterparts in 5-HT_3A_ L1-MX in the biogenesis of α7 nACh and 5-HT_3A_ receptors.

α7 nAChR has long been pursued as a drug target for cognitive dysfunction in schizophrenia and Alzheimer’s disease. Drug development approaches have focused on developing compounds that target the primary ligand binding site (orthosteric site) at the ECD, or allosteric sites located in the TMD. Despite promising preclinical data and encouraging early-phase trials, many compounds, such as encenicline (EVP-6124), bradanicline (TC-5619), and DMXB (GTS-21), have failed to meet significant endpoints in larger late-phase trials. One reason for the failure are off-target effects arising from conserved structures in ECDs and TMDs across members of the pLGIC subfamily (Walling et al., 2016; Bertrand and Terry, 2018; Papke and Horenstein, 2021). To identify structural elements that provide selectivity and can serve as potential novel drug targets, we have investigated the ICD of the pLGIC superfamily, the region that is most variable in length and amino acid composition (Jansen et al., 2008; Goyal et al., 2011; Mnatsakanyan et al., 2015; Pandhare et al., 2016; Pirayesh et al., 2020; Stuebler and Jansen, 2020; Do and Jansen, 2023). Our contrasting findings on conserved residues in the L1-MX fragments of different pLGIC members, α7 nAChR vs. 5-HT_3A_ receptor, indicate the existence of distinct molecular mechanisms associated with the L1-MX fragment in assisting the biogenesis of the receptors. Our findings here, thus, position the L1-MX fragment as a potential target for developing new therapeutics that can minimize adverse effects.

A previous study demonstrated that a single Ala replacement at L336A impairs the surface expression of α7 nAChRs as measured by binding of ^125^I-αBgt in intact HEK cells (Mukherjee et al., 2009). This previous study also demonstrates Ala-replacements at other residues in L1-MX, L335, N337, and W338 (Fig. 2 A) impair the surface expression of α7 nAChRs in HEK cells. In the muscle δ nACh subunit, K351 corresponds to R332 of α7 nACh subunit. The ubiquitination at this Lys residue leads to the retention of receptors in the Golgi apparatus. Replacement of the Lys with Arg, K351R, reduces ubiquitination, and the surface expression of the chimeric muscle nAChR receptors is increased (Rudell et al., 2014; Rudell et al., 2020). Likely, residues L336 and R332 are positively involved in the assembly and trafficking of nAChRs.

However, in α7 nAChR, when L336 and R332 were preserved in our single Ala replacement construct, MX-A or W330A, α7 current was still abolished. Interestingly, our co-expression of the MX-A construct with chaperone RIC-3 or NACHO could rescue some α7 current (Fig. 3 D And Fig. 4). When both L336 and R332 were replaced by Ala residues, and W330 was kept intact, α7 current was also abolished; and the presence of RIC-3 or NACHO could barely rescue α7 current. Taken together, our data, supported by the previous studies, demonstrate that the MX segment or subdomain is crucial for the functional assembly and surface expression of α7 nAChR. Our finding also provides an approach to turn on and off the α7 current, which we hope will be beneficial for future therapeutic development targeting α7 nAChR.

Previously, it was shown that L264 and G265 in the M2 segment of α7 nAChR are crucial for NACHO-mediated assembly and trafficking. However, this study did not establish a binding site for NACHO (Kweon et al., 2020). Our pull-down data indicate that NACHO strongly binds to α7 L1-MX peptide containing 25 amino acids. Our data also demonstrate that in co-expression with NACHO, α7 currents are rescued in constructs containing either W330A or L336A and R332A. Taken together, our findings clearly demonstrate that NACHO binds the L1-MX segment of the ICD domain of the α7 nACh subunit. To the best of our knowledge, this is the first time, a binding location for NACHO has been identified in α7 nAChR and also in any pLGIC.

α7 nAChR is expressed in neurons, astrocytes, immune cells, and multiple cancer cell lines (Sargent, 1993; Dani and Bertrand, 2007; Wessler and Kirkpatrick, 2008). Recent data provide cumulative evidence that α7 nAChR mediates cell proliferation, angiogenesis, and anti-apoptosis, making it a potential drug target in cancer therapy or prevention (Zia et al., 2000; Cucina et al., 2012; Gergalova et al., 2012; Pillai and Chellappan, 2012; Schuller, 2012; Onder Narin et al., 2021). Further studies are recommended to determine whether NACHO promotes the assembly and stabilization of α7 nAChR in cancer cell lines.

## CONCLUSION

Here, we report a small peptide of 25 amino acids, derived from the L1-MX segment of α7 nAChR’s ICD domain, to strongly bind NACHO. Single (W330A), double (R332A and L336A), or triple (W330A, R332A, and L336A) Ala replacements at conserved sites in the L1-MX segment of full-length α7 nAChR abolished α7 current. However, in co-expression with NACHO, the α7 currents are rescued from constructs containing W330A or R332A and L336A. In addition, we demonstrate that NACHO stabilizes α7 pentamer via non-covalent interactions. RIC-3 also binds to this 25-amino-acid peptide as it does with synthetic 5-HT_3A_ peptide derived from the L1-MX segment of 5-HT_3A_ receptor as shown in our previous study (Do and Jansen, 2023).

Similar changes in side chains of conserved residues in this L1-MX chaperone binding segment abrogate α7 current and reduce 5-HT_3A_ current presenting a critical role of the L1-MX segment in the biogenesis of α7 nACh and 5-HT_3A_ receptors. Overall, this segment is a crucial hotspot for the biogenesis, and chaperone interaction. It therefore presents a promising site for additional investigations, including as a potential drug target.

## Data Availability

Western blot data for biological or technical replicates, as well as the original two-electrode voltage clamp recording files are available from the corresponding authors upon reasonable request.

## Acknowledgment

Research reported in this publication was supported by the National Institute of Neurological Disorders and Stroke of the National Institutes of Health under award number R01NS077114 (to M.J.). We thank Dr. Lindstrom (University of Pennsylvania, Philadelphia, PA) and Dr. Treinin (Hebrew University, Jerusalem, Israel) for the plasmid gifts.

## Author contributions

QHD and MJ conceived the project and designed the experimental procedures. QHD, IKC, and PNG acquired the data. JJT, and RR assisted in data acquisition. QHD and MJ analyzed, discussed, and interpreted the data. QHD and MJ and wrote the manuscript. All authors contributed to the final version of the manuscript.

